# Membrane Nanoparticles Derived from ACE2-rich Cells Block SARS-CoV-2 Infection

**DOI:** 10.1101/2020.08.12.247338

**Authors:** Cheng Wang, Shaobo Wang, Yin Chen, Jianqi Zhao, Songling Han, Gaomei Zhao, Jing Kang, Yong Liu, Liting Wang, Xiaoyang Wang, Yang Xu, Song Wang, Yi Huang, Junping Wang, Jinghong Zhao

## Abstract

The ongoing COVID-19 epidemic worldwide necessitates the development of novel effective agents against SARS-CoV-2. ACE2 is the main receptor of SARS-CoV-2 S1 protein and mediates viral entry into host cells. Herein, the membrane nanoparticles prepared from ACE2-rich cells are discovered with potent capacity to block SARS-CoV-2 infection. The membrane of human embryonic kidney-239T cell highly expressing ACE2 is screened to prepare nanoparticles. The nanomaterial termed HEK-293T-hACE2 NPs contains 265.1 ng mg^−1^ of ACE2 on the surface and acts as a bait to trap SARS-CoV-2 S1 in a dose-dependent manner, resulting in reduced recruitment of the viral ligand to host cells. Interestingly, SARS-CoV-2 S1 can translocate to the cytoplasm and affect the cell metabolism, which is also inhibited by HEK-293T-hACE2 NPs. Further studies reveal that HEK-293T-hACE2 NPs can efficiently suppress SARS-CoV-2 S pseudovirions entry into human proximal tubular cells and block viral infection with a low half maximal inhibitory concentration. Additionally, this biocompatible membrane nanomaterial is sufficient to block the adherence of SARS-CoV-2 D614G-S1 mutant to sensitive cells. Our study demonstrates a easy-to-acheive memrbane nano-antagonist for curbing SARS-CoV-2, which enriches the existing antiviral arsenal and provides new possibilities to treat COVID-19.

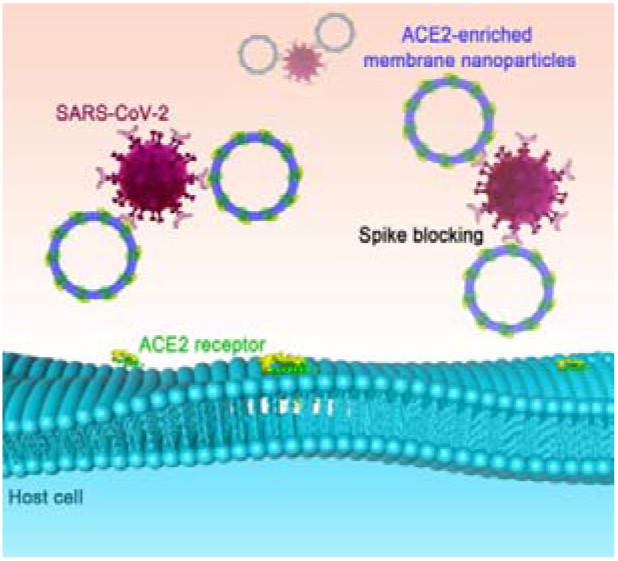
Graphical Table of Contents

---

The outbreak of corona virus disease 2019 (COVID-19) caused by severe acute respiratory syndrome coronavirus 2 (SARS-CoV-2) has infected over 20,000,000 individuals worldwide, resulting in more than 730,000 deaths.^1^ Fever, cough, and myalgia or fatigue are the common symptoms of COVID-19 patients.^2^ Once the illness worsens, acute respiratory distress syndrome and respiratory failure, sepsis, and acute kidney injury are the common lethal complications.^3^ In addition to the supportive and symptomatic care, development of therapies that target the viral pathogenic processes including cellular association, membrane penetration, endosomal escape, viron uncoating, and genome replication are instrumental to improve the therapeutic outcome.^4^

The attachment of SARS-CoV-2 to cell membrane is the initial step of the pathogenesis of COVID-19, in which the viral spike (S) protein that can be divided into S1 and S2 subunits after degradation by protease is attributable.^5^ S1 is responsible for recognizing host receptors, while S2 mediates viral fusion into the cytoplasm.^6^ Angiotensin-converting enzyme-2 (ACE2), a negative regulator of the renin-angiotensin system, is abundant in kidney, lung, and intestine.^7^ SARS-CoV-2 S1 binds to ACE2 at a high affinity of 8.02 nM.^8^ The cells insensitive to SARS-CoV-2 turn to be susceptible to the virus after transfection with ACE2. Additionally, anti-ACE2 serum and human intestinal defensin able to cloak ACE2 we recently reported were efficient to inhibit SARS-CoV-2 invasion,^6, 8^ indicating that ACE2 is a critical receptor for SARS-CoV-2.

Similar to that the extracellular part of the coxsackievirus and adenovirus receptor is used to inhibit Coxsackie-B-viruses,^9^ recombinant human ACE2 (rhACE2) at clinical grade markedly reduces the early infection of SARS-CoV-2 and protects human blood vessel and kidney organoids.^10^ Due to the natural location of ACE2 on the cell membrane, and because membrane with active ingredients are potential drug candidates,^11^ we attempt to apply the membrane of human cells abundant of ACE2 to cope with SARS-CoV-2. Cell membrane-based nanoparticles (CMBNPs) are designed to overcome the shortcoming of the uneven size of membranes. Taking the advantage of functional elements on human platelet membranes, we have developed some CMBNPs with the capabilities of tumor targeting and immune escape.^12–13^

In this study, we selected the membrane of human embryonic kidney-239T cells highly expressing human ACE2 (HEK-293T-hACE2) to prepare CMBNPs after analyzing the ACE2 content in five human cells (Figure 1). The antiviral activity and mechanism of action of this nanomaterial were investigated by pseudovirus neutralization assay, spike adhesion experiment, and proteomics analysis. Our study demonstrates an efficient nano-antagonist that can be easily prepared in general laboratories against SARS-CoV-2, which is a feasible solution to resolve the shortage of effective measures to treat COVID-19.

**Figure 1.**
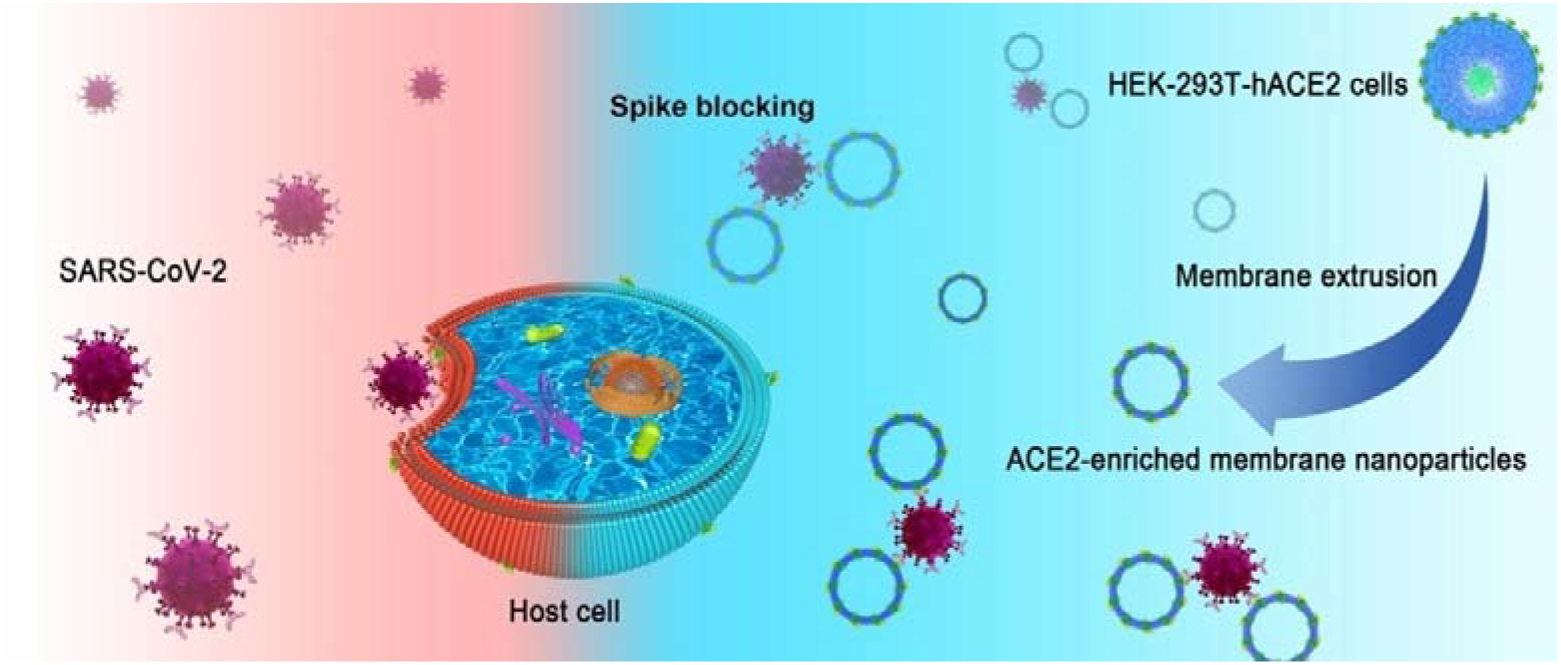
Diagrammatic drawing depicting the preparation and function of HEK-293T-hACE2 NPs.

## RESULTS AND DISCUSSION

### Preparation and characterization of HEK-293T-hACE2 NPs

ACE2 is rich in renal tubular epithelium, lung alveolar epithelial cells, and enterocytes of the small intestine, as revealed by immunohistochemistry and transcriptomics and proteomics analysis.^7, 14^ We collected five cells including HEK-293T, HEK-293T-hACE2, human proximal tubular cell HK-2, Caco-2 enterocytes, and A549 type II pneumocytes and determined their ACE2 content by Western blot. Owing to the codon optimization, HEK-293T-hACE2 was superior to HEK-293T, Caco-2, HK-2, and A549 cells at carrying the viral receptor (Figure 2a). Immunofluorescence observed that ACE2 was mostly located on the cell membrane (Figure 2b). HEK-293T-hACE2 cells were then selected and processed by repeated freezing and thawing to separate the membrane, which was broken by sonication and applied to fabricate CMBNPs using a classical extrusion method.^15^ Transmission electron microscope (TEM) analysis revealed that similar to red blood cells (RBCs)-derived NPs prepared using the same size of polycarbonate membrane,^16^ HEK-293T-hACE2 NPs were approximately 100 nm with a preferable dispersity in solution (Figure 2c). The diameter of this negatively charged nanomaterial detected by dynamic light scattering (DLS) averaged 169 nm (Figure 2d). HEK-293T NPs prepared using the same method had comparable ζ-potential and diameter with HEK-293T-hACE2 NPs.

**Figure 2.**
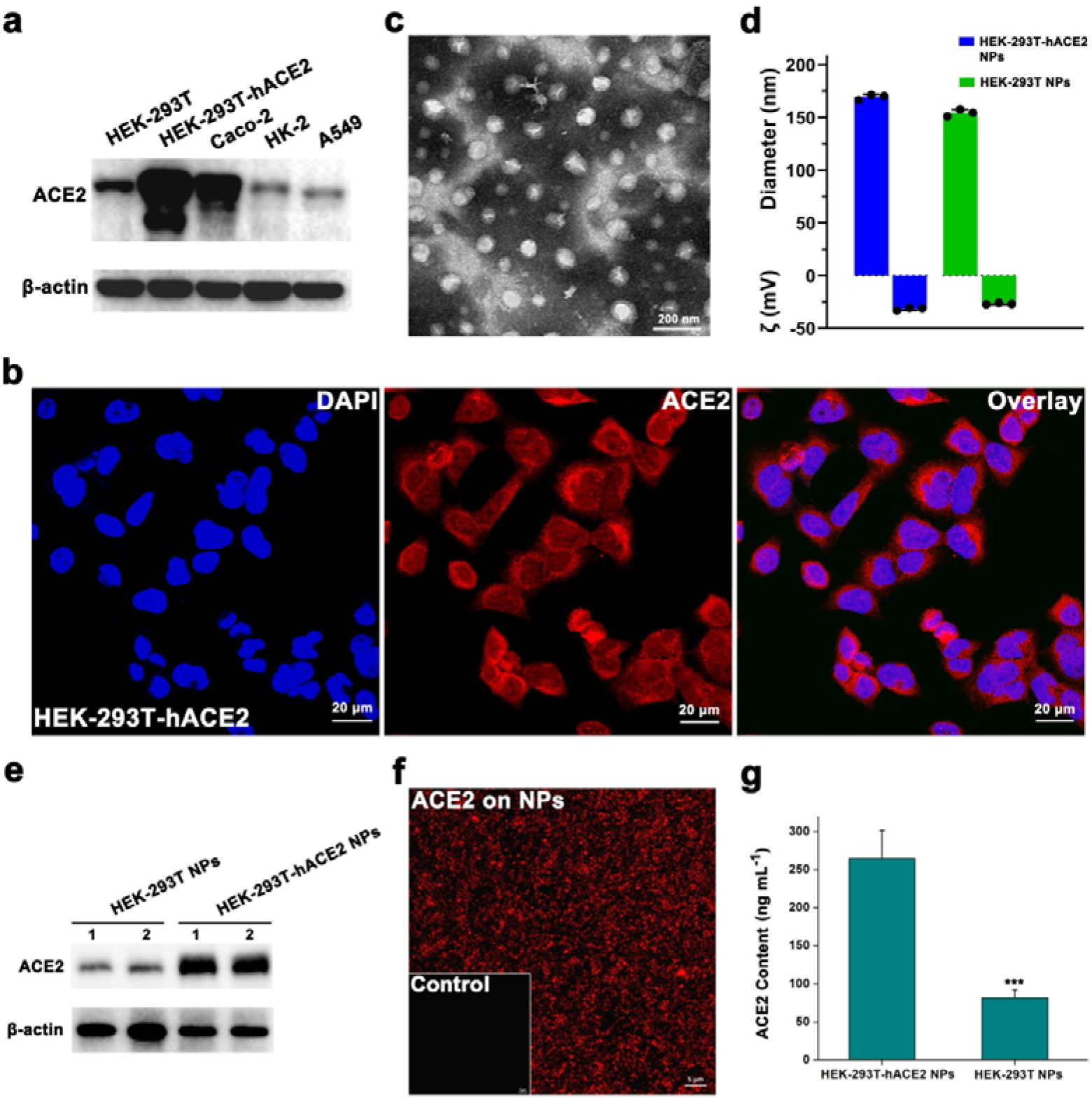
ACE2 screening in human cells and the characterization of HEK-293T-hACE2 NPs. (a) Western blot determining the ACE2 contents in five human cells. β-actin is the reference. (b) Immunofluorescence microscopy showing the location of ACE2 (red) in HEK-293T-hACE2 cells. Nuclei are stained by DAPI (blue). Scale bar indicates 20 μm. (c) TEM image of HEK-293T-hACE2 NPs. The scale is 200 nm. (d) Hydrodynamic diameters and surface charges of the membrane nanomaterials. The results are shown as the means ± SD. (e) Immunobloting determining the ACE2 contents in nanomaterials. β-actin is the reference. (f) Immunofluorescence microscopy observing the abundence of ACE2 (red) on HEK-293T-hACE2 NPs. Scale bar indicates 5 μm. (g) ELISA measuring the ACE2 contents in nanomaterials. ***, *P* < 0.001.

### Inhibition of HEK-293T-hACE2 NPs on SARS-CoV-2 S1

SARS-CoV-2 S1 containing a receptor-binding domain (RBD) is the ligand of ACE2.^17^ HEK-293T-hACE2 NPs carried abundant ACE2 (Figure 2e and 2f). Enzyme-linked immunosorbent assay (ELISA) determined that the content of ACE2 in HEK-293T-hACE2 NPs was 265.1 ng mg^−1^, 3.2-fold higher than that in HEK-293T NPs (Figure 2g, *P* < 0.001). To evaluate the bioactivity of HEK-293T-hACE2 NPs, we initially immobilized biotinylated SARS-CoV-2 S1-RBD on Sartorius streptavidin (SA) biosensors and determined the recruitment of CMBNPs by biolayer interferometry (BLI). In line with the discrepancy in ACE2 loading, more HEK-293T-hACE2 NPs coated SARS-CoV-2 S1-RBD than HEK-293T NPs (Figure 3a). As SARS-CoV-2 S1 adheres to the surface of sensitive cells including HK-2,^8^ we further performed a cell experiment, in which HK-2 cells were exposed to 10 μg mL^−1^ of SARS-CoV-2 S1.

**Figure 3.**
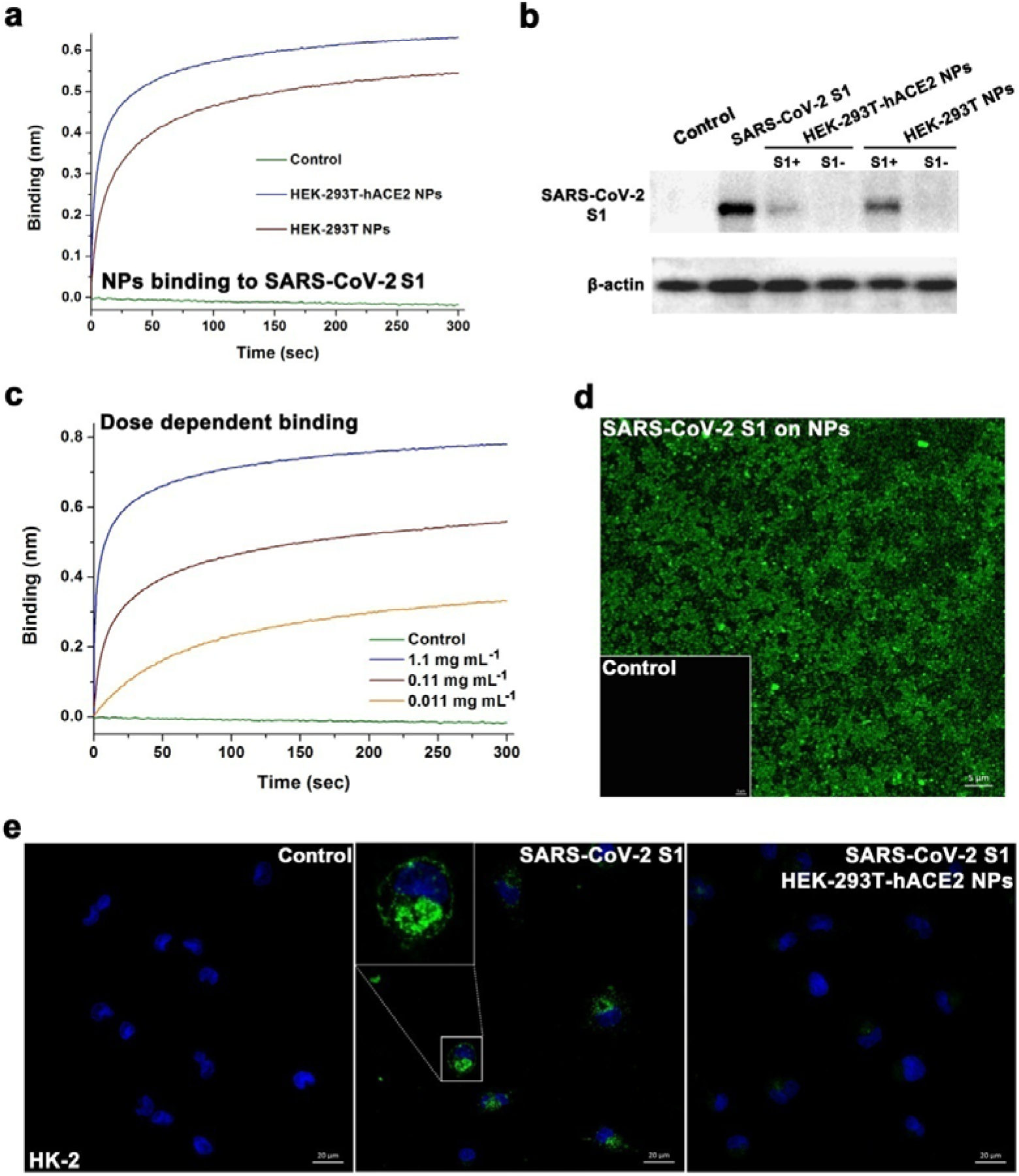
Inhibition of HEK-293T-hACE2 NPs on SARS-CoV-2 S1. (a) Binding kinetics for NPs and SARS-CoV-2 S1-RBD loaded on SA biosensors. HEK-293T-hACE2 NPs and HEK-293T NPs are 1.1 mg mL^−1^ in PBS. (b) Western blot determining the content of S1 binding to HK-2 in the absence and presence of NPs. β-actin is the reference. (c) Binding kinetics for increasing concentrations of HEK-293T-hACE2 NPs and SARS-CoV-2 S1-RBD loaded on SA biosensors. (d) Immunofluorescence microscopy observing the adherence of SARS-CoV-2 S1 (green) on HEK-293T-hACE2 NPs. Scale bar indicates 5 μm. (e) Immunofluorescence microscopy revealing the protection of HEK-293T-hACE2 NPs on HK-2 cells exposed to SARS-CoV-2 S1 (green). The region of interest in SARS-CoV-2 S1-treated group is magnified in the embedding graph. Nuclei are stained using DAPI (blue). Scale bar indicates 20 μm.

Western blot detected that HK-2 recruited plenty of SARS-CoV-2 S1 without treatment (Figure 3b). Nevertheless, when the viral protein was pretreated with 2.5 mg mL^−1^ of CMBNPs (based on the membrane proteins), the recruitment of SARS-CoV-2 S1 was dramatically decreased. Notably, HEK-293T-hACE2 NPs was more efficient than HEK-293T NPs at neutralizing SARS-CoV-2 S1. Binding assay further revealed that HEK-293T-hACE2 NPs interacted with SARS-CoV-2 S1-RBD in a dose-dependent manner (Figure 3c), consistent with the results obtained by immunoblotting (Figure S1). Immunofluorescence observed that SARS-CoV-2 S1 was adsorbed on HEK-293T-hACE2 NPs (Figure 3d) and the protein adherence was blocked by human defensin 5 (HD5, Figure S2), an endogenous lectin-like peptide capable of binding to and cloaking ACE2.^8^ Accordingly, few viral proteins adhered to HK-2 cells after pretreatment with HEK-293T-hACE2 NPs (Figure 3e).

Intriguingly, SARS-CoV-2 S1 not only located on the cell membrane but in the cytoplasm of HK-2, in line with the translocation of spike in Caco-2 cells.^8^ Recent multi-level proteomics analysis showed that individual proteins of SARS-CoV-2 reshaped the central pathways of host cells, causing metabolic disorders including the dysfunctions in mitochondrial translational termination and cholesterol metabolic abnormalities.^18^ SARS-CoV-2 S1 might has a potential effect on cellular metabolism we speculated beyond receptor recognition. A tandem mass tags (TMT) proteomics research was then conducted to verify the conjecture. The contents of fifty-two proteins (Table S1) including voltage-dependent anion channel (VDAC) 1, VDAC2, and VDAC3 were significantly altered (fold-change > 1.2) in HK-2 cells after exposure to 100 ng mL^−1^ of SARS-CoV-2 S1 for 48 h (Figure 4a). Kyoto encyclopedia of genes and genomes (KEGG) enrichment analysis discovered six signaling pathways (Table S2), i.e., cGMP-PKG signaling pathway, NOD-like receptor signaling pathway, calcium signaling pathway, cholesterol metabolism, necroptosis, and ferroptosis, which were influenced by SARS-CoV-2 S1 (Figure 4b).

**Figure 4.**
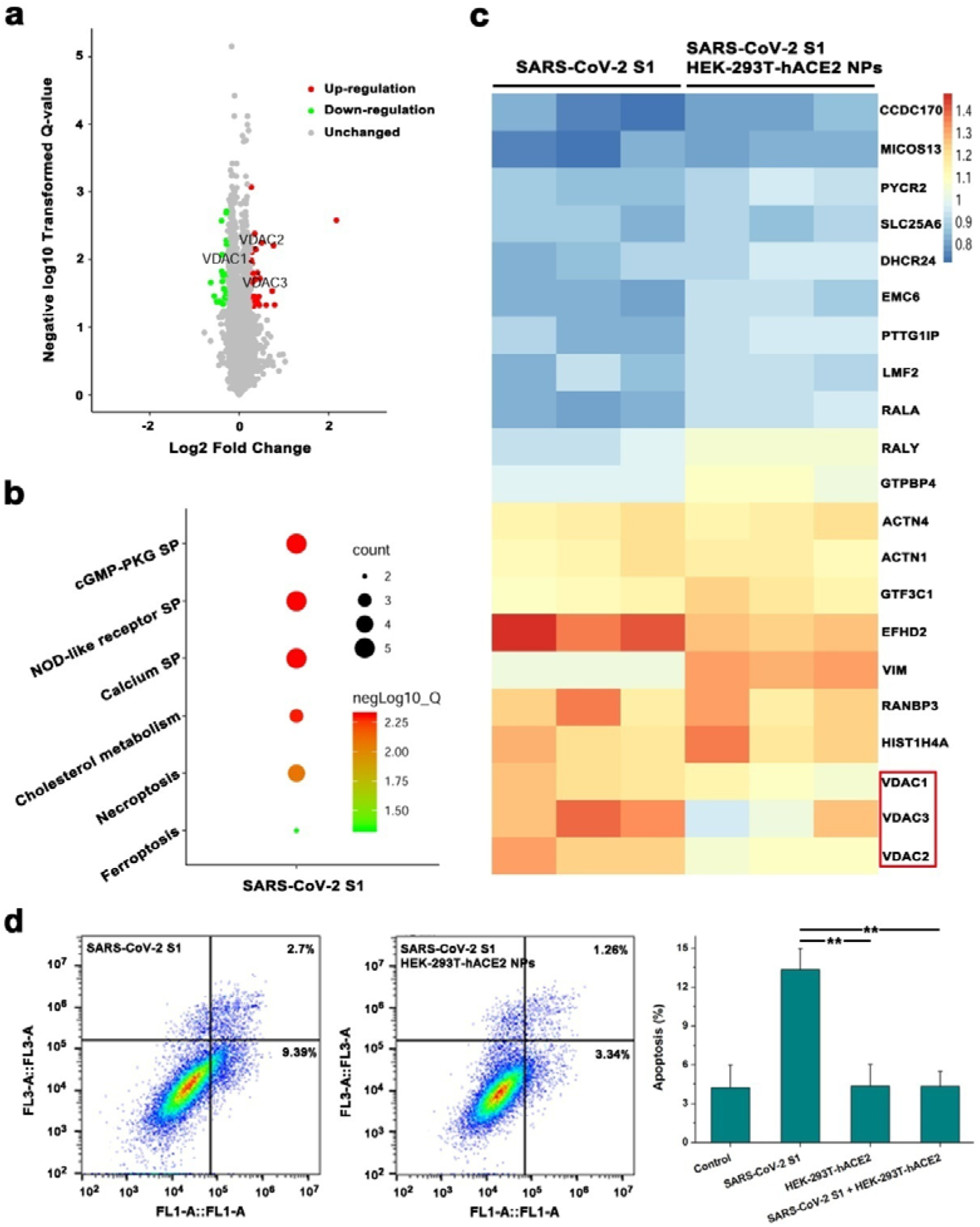
Alleviation of HEK-293T-hACE2 NPs to the metabolic disturbance of HK-2 cells exposed to SARS-CoV-2 S1. (a) Volcano plot showing the changes of cell proteins. The increased proteins VDAC1-3 are labeled. (b) Altered signaling pathways (SPs) in HK-2 exposed to SARS-CoV-2 S1. (c) Heat map displaying the correction of HEK-293T-hACE2 NPs to the imbalanced cell proteins induced by SARS-CoV-2 S1. VDAC1-3 are highlighted in a red frame. (d) Flow cytometry of HK-2 treated with SARS-CoV-2 S1 in the absence and presence of HEK-293T-hACE2 NPs. **, *P* < 0.01.

The up-regulation of VDAC1-3, three mitochondrial membrane porins involved in bioenergetic failure and cell apoptosis,^19–20^ were attributable. In the presence of HEK-293T-hACE2 NPs, the imbalance of VDAC1-3 was corrected (Figure 4c), and the number of changed proteins was reduced to thirty-one (Table S3), by which no signaling pathway was enriched by KEGG yet. Flow cytometry revealed that 100 ng mL^−1^ of SARS-CoV-2 S1 increased the apoptisis of HK-2, which was significantly inhibited by 100 μg mL^−1^ of HEK-293T-hACE2 NPs (Figure 4d), supporting that HEK-293T-hACE2 NPs treatment alleviates the effect of SARS-CoV-2 S1 on the metabolic process of host cells.

### Suppression of HEK-293T-hACE2 NPs on SARS-CoV-2 D614G-S1

Along with the viral mutation, a variant SARS-CoV-2 S1 with the Asp^614^ replaced with Gly (D614G-S1) is globally prevalent.^21^ The single site mutation makes SARS-CoV-2 three to nine-fold more infectious than the wild type.^22^ We analyzed the molecular interaction by BLI and found that D614G-S1 bound to ACE2 at 13.3 nM (Figure 5a), comparable to the affinity of SARS-CoV-2 S1 binding to ACE2,^8^ which sustained that the stronger contagiosity of D614G-SARS-CoV-2 than its ancestral form is not due to the enhanced receptor recruitment.^23^ Similarly, D614G mutation did not alter the dose-dependent bindings of HEK-293T-hACE2 NPs to SARS-CoV-2 S1 (Figure 5b). Immunofluorescence observed that HEK-293T-hACE2 NPs markedly lowered the adherence of D614G-S1 to HK-2 cells (Figure 5c), indicative of the efficacy of this membrane nanomaterial at blocking mutant SARS-CoV-2.

**Figure 5.**
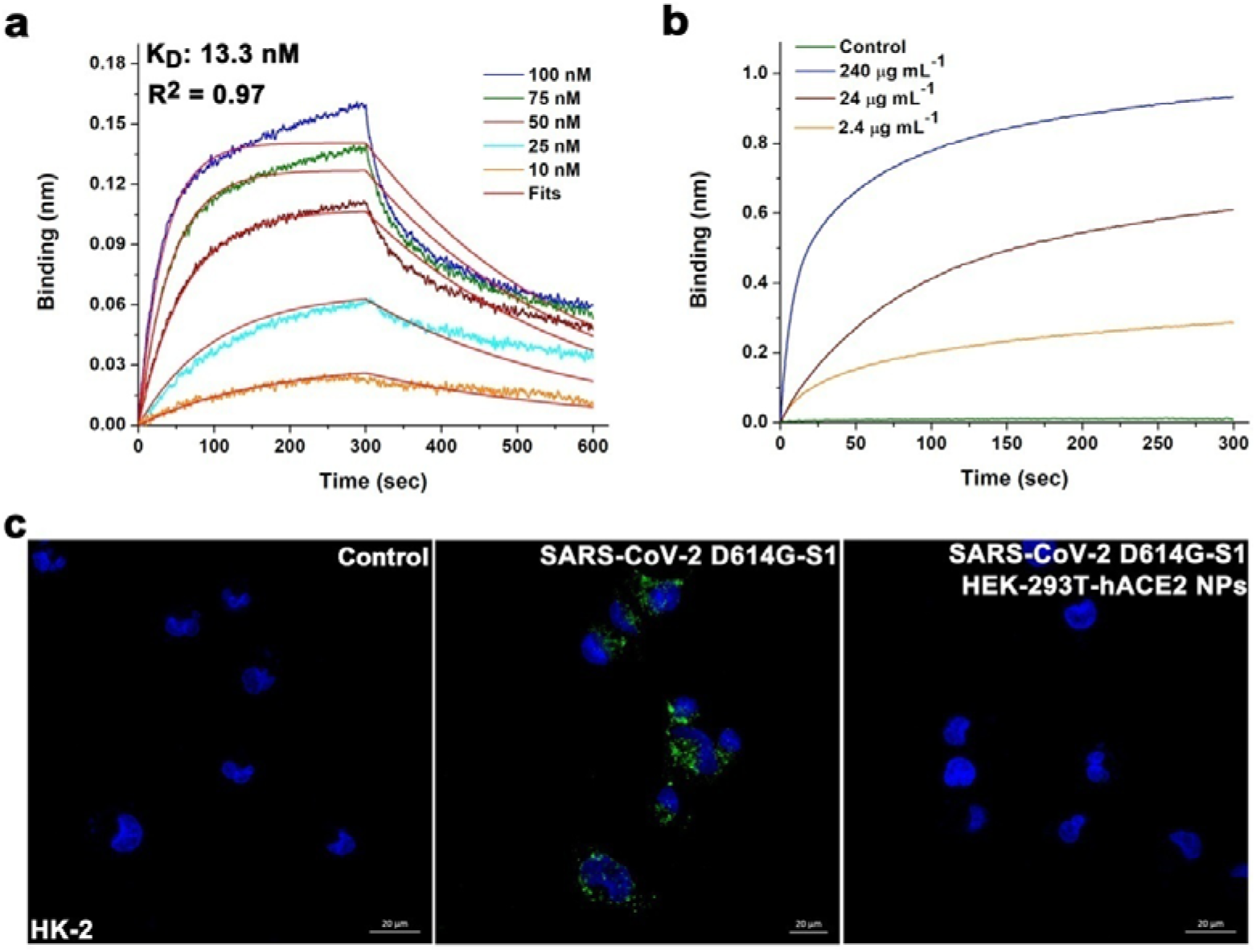
Inhibition of HEK-293T-hACE2 NPs on SARS-CoV-2 D614G-S1. (a) Binding kinetics for ACE2 and D614G-S1 loaded on SA biosensors. The fitting curves are in red. (b) Binding kinetics for increasing concentrations of HEK-293T-hACE2 NPs and D614G-S1 loaded on SA biosensors. (c) Immunofluorescence microscopy revealing the protection of HEK-293T-hACE2 NPs on HK-2 cells exposed to D614G-S1 (green). Nuclei are stained using DAPI (blue). Scale bar indicates 20 μm.

### Antiviral action of HEK-293T-hACE2 NPs

For insights into the antiviral activity of HEK-293T-hACE2 NPs, we employed SARS-CoV-2 S pseudovirions containing psPAX2 and luciferase reporter system to infect HK-2 cells. SARS-CoV-2 S1 binds to ACE2 and enters host cells after the proteolysis of transmembrane protease serine 2 (TMPRSS2) at the S1/S2 boundary.^24^ The intracellular proprotein convertase furin also promotes SARS-CoV-2 entry by pre-activating viral spike.^25^ Because human kidney is abundant of ACE2,^26^ TMPRSS2, and furin (Figure S3), SARS-CoV-2 S pseudovirions were largely endocytosed by HK-2 cells after a 1 h of co-incubation (Figure 6a). HEK-293T-hACE2 NPs treatment markedly reduced viral entry, in keeping with its inhibiting effect on SARS-CoV-2 S1. TEM explained that pseudovirions were adsorbed on HEK-293T-hACE2 NPs (Figure 6b), forming a coronavirus-like complex.^27^ Furthermore, luciferase assay was carried out 48 h post the infection of SARS-CoV-2 S pseudovirions. HEK-293T-hACE2 NPs exerted a dose-dependent antiviral activity (Figure 6c). The half maximal inhibitory concentration (IC_50_) of HEK-293T-hACE2 NPs was 431.2 μg mL^−1^, lower than that of human lung epithelial type II cells-derived NPs.^28^ According to the result shown in Figure 2g, HEK-293T-hACE2 NPs at the concentration of IC_50_ contained 0.114 μg mL^−1^ of ACE2, comparable to the IC_50_ of rhACE2 (0.1 μg mL^−1^) at blocking SARS-CoV-2.^29^

**Figure 6.**
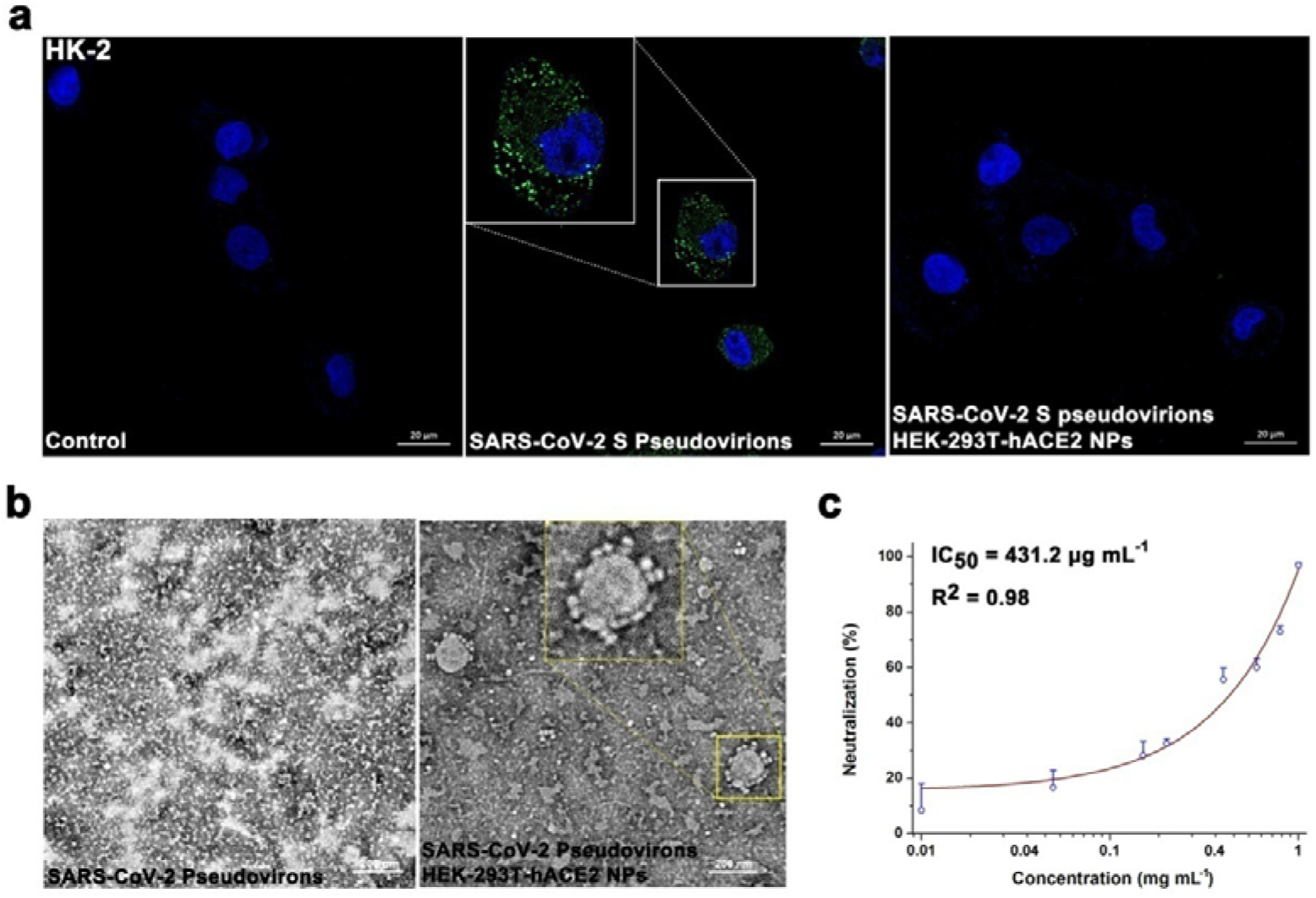
Antiviral evaluation of HEK-293T-hACE2 NPs. (a) HEK-293T-hACE2 NPs protect HK-2 cells from SARS-CoV-2 S pseudovirons invasion, as revealed by S1 immunofluorescence (green). Nuclei are stained by DAPI (blue). Scale bar indicates 20 μm. The region of interest in pseudovirons-treated group is magnified in the embedding graph. (b) TEM image of SARS-CoV-2 S pseudovirons adsorbed on HEK-293T-hACE2 NPs. The region of interest is magnified in the embedding graph. The scale bar is 200 nm. (c) Dose-dependent antiviral action of HEK-293T-hACE2 NPs. The fitting curve is in wine.

### Biocompatibility of HEK-293T-hACE2 NPs

Overexpression of membrane receptors does not affect the biocompatibility of human CMBNPs.^30^ We incubated increasing concentrations of HEK-293T-hACE2 NPs with human umbilical vein endothelial cells (HUVECs) for 24 h. The material did not influence the cell survival at doses up to 500 μg mL^−1^ (based on the membrane proteins, Figure 7a). The negligible hemolysis additionally indicated the biosafety of this material *in vitro* (Figure 7b). Animal experiments found that intravenous injections of 25 mg kg^−1^ of HEK-293T-hACE2 NPs had less of effect on the counts of RBCs, white blood cells (WBCs), and platelets (PLTs, Figure 7c) or the hemoglobin content (Figure S4) in the blood. The major organs including heart, liver, spleen, lung, and kidney were collected at seven days post-initial-injection. Hematoxylin and eosin (HE) staining observed no pathological changes in the tissues (Figure 7d), suggestive of the nontoxicity *in vivo*, thus laying a good foundation for the application of HEK-293T-hACE2 NPs as a nano-antagonist against SARS-CoV-2.

**Figure 7.**
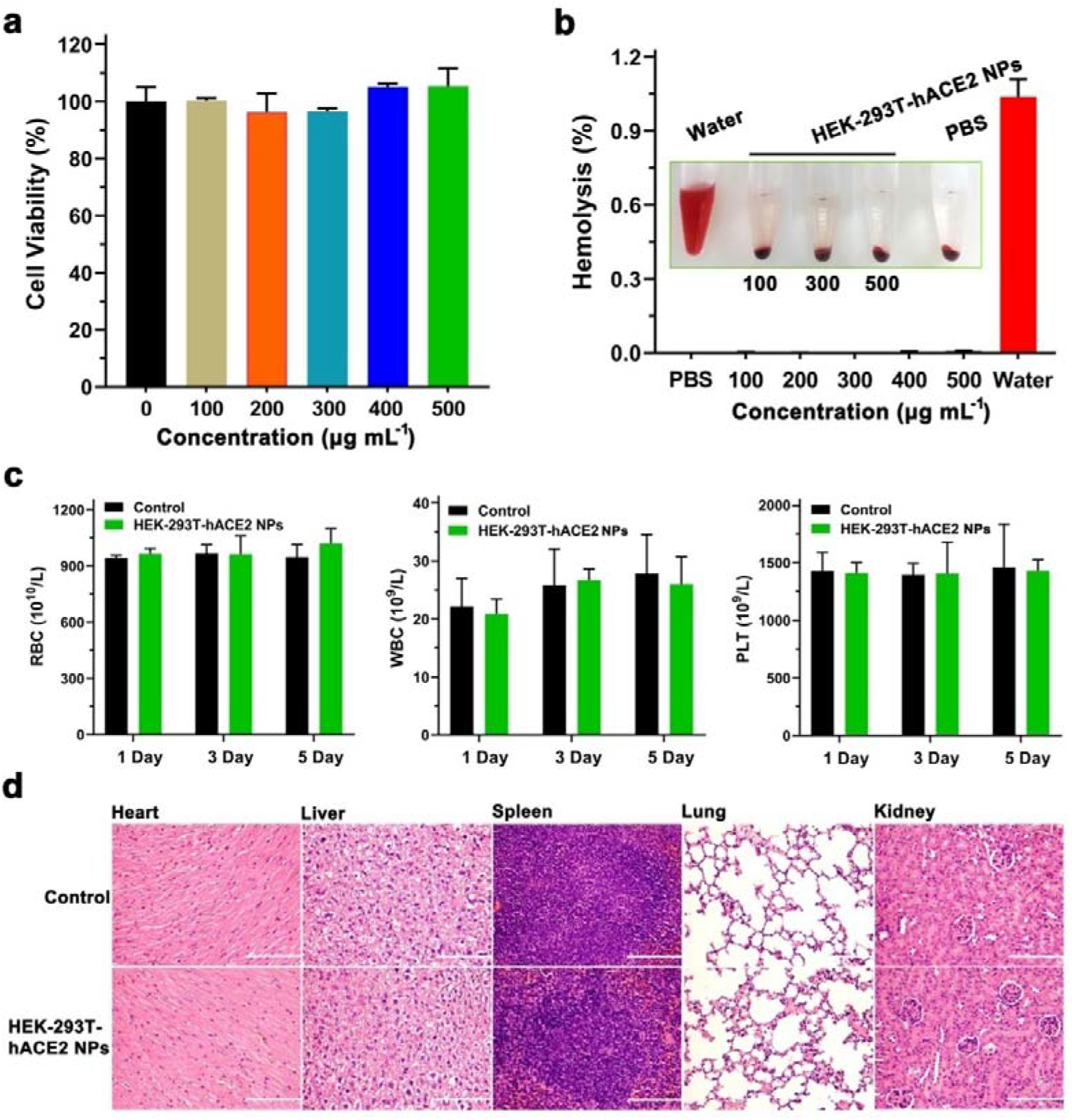
Toxicity evaluation of HEK-293T-hACE2 NPs. (a) The survival of HUVECs exposed to different concentrations of HEK-293T-hACE2 NPs for 24 h. The results are shown as the means ± SD. (b) Hemolysis of different doses of HEK-293T-hACE2 NPs. (c) Counts of RBC, WBC, and PLT in mouse blood at 1, 3, and 5 days post-injection. The results are presented as the means ± SD. (d) HE staining of the organs of mice treated with NaCl and HEK-293T-hACE2 NPs. The scale is 200 μm.

## CONCLUSION

A membrane nanomaterial derived from HEK-293T-hACE2 cells against SARS-CoV-2 was prepared and evaluated in the present study. HEK-293T-hACE2 NPs contained abundant ACE2, a critical receptor of SARS-CoV-2 S1. Due to the competitive inhibition, HEK-293T-hACE2 NPs bound to SARS-CoV-2 S1 and blocked the viral ligand adhering to human renal tubular epithelial cells in a dose-dependent manner. The effect of SARS-CoV-2 S1 on cellular metabolism was suppressed by HEK-293T-hACE2 NPs as well. Owing to the spike recruitment, HEK-293T-hACE2 NPs adsorbed SARS-CoV-2 S pseudovirons on the material surface and blocked viral entry into the cytoplasm, thus endowing inspiring pretections for host cells from the viral infection. As there is a shortage of effective measures to treat COVID-19, the biocompatible and easy-to-acheive nano-antagonist we think may be a useful therapeutic candidate for curbing SARS-CoV-2.

## EXPERIMENTAL SECTION

### Western Blot

HEK-293T, Caco-2, HK-2, A549 cells were obtained from the cell bank of Chinese Academy of Sciences (CAS, Shanghai, CHN). HEK-293T-hACE2 cells were obtained from Prof. Lilin Ye.^31^ The cells were cultured in dulbecco’s modified eagle medium (DMEM, 11995065, Gibco, Thermo Fisher Scientific, Shanghai, CHN) containing 10% fetal bovine serum (FBS, 10100147, Gibco). Geneticin (400 μg mL^−1^, ST081, Beyotime, Shanghai, CHN) was added to the culture medium of HEK-293T-hACE2 for stress screening. After three times of wash with sterile PBS, cells were collected and lysed for protein extraction. A total of 25 μg of each protein sample is resolved by 10% SDS-PAGE. A primary anti-ACE2 rabbit polyclonal antibody (1:1000, 10108-T26, Sino Biological, Beijing, CHN) and a goat anti-rabbit secondary antibody (1:1000, A0208, Beyotime) were employed to detect ACE2. β-actin determined by a mouse monoclonal antibody (AA128, Beyotime, 1:1000) was a reference.

To determine the content of SARS-CoV-2 S1 adhered to HK-2, cells were seeded into a 6-well plate at a density of 1 × 10^6^ cells/well. Recombinant SARS-CoV-2 S1 (20 μg mL^−1^, 40591-V08H, Sino Biological) containing a His-tag preincubated with CMBNPs at 37 °C for 1 h were added to HK-2 cells and incubated for another 1 h. After three times of wash with PBS, cells were collected and lysed. A primary anti-His-tag mouse monoclonal antibody (1:1000, AF5060, Beyotime) and a goat anti-mouse secondary antibody (1:1000, A0216, Beyotime) were employed to detect SARS-CoV-2 S1. These experiments were repeated three times in different days.

### Immunofluorescence Microscopy

HEK-293T-hACE2 and HK-2 cells were seeded into 12-well plates with sterile glass slides at a density of 2 × 10^5^ cells/well. A total of 100 μL of HEK-293T-hACE2 NPs (2.5 mg mL^−1^, based on the membrane proteins) co-incubated with or without 10 μg mL^−1^ of SARS-CoV-2 S1 at 37 °C for 1 h was dripped on sterile glass slides coated with polylysine. To observe the adherences of SARS-CoV-2 S1, D614G-S1 (40591-V08H3, Sino Biological), and pseudovirons to host cells, HK-2 was exposed to 10 μg mL^−1^ of spike or 1.98 × 10^7^ TU mL^−1^ of SARS-CoV-2 S pseudovirons (Genewiz, Suzhou, Jiangsu Province, CHN) for 1 h. Cells cultured overnight and HEK-293T-hACE2 NPs were washed with PBS and fixed in 4% paraformaldehyde. A primary anti-ACE2 rabbit polyclonal antibody (1:200, 10108-T60, Sino Biological) and a goat anti-rabbit secondary antibody (1:500, A0516, Beyotime) were employed to stain ACE2. A primary anti-spike rabbit monoclonal antibody (1:100, 40150-R007, Sino Biological) and a goat anti-rabbit secondary antibody (Alexa Fluor 488, Invitrogen, Thermo Fisher Scientific) were used to stain S1. Nuclei were labelled with 2-(4-amidinophenyl)-6-indolecarbamidine dihydrochloride (DAPI, C1002, Beyotime). A Zeiss LSM 780 NLO confocal microscope was applied to observe the cells.

### Preparation of HEK-293T-hACE2 and HEK-293T NPs

HEK-293T-hACE2 cells collected by trypsinization and centrifugation at 1500 rpm for 5 min were washed with cold sterile PBS and frozen at −80 °C and thawed at room temperature (repeated three times). The cracked membrane was obtained by centrifugation at 8,000 rpm for 10 min, washed with cold PBS containing protease inhibitors and sonicated with a Sonics (Newtown, CT, US) Vibra-Cell VCX-500 ultrasonic processor for 10 min at a power of 120 W. CMBNPs were prepared by continuously extruding the cell membrane for 13 times using a LiposoFast-Basic mini extruder (Avanti Polar Lipids, Alabaster, AL, US) equipped with a 200 nm porous membrane.^15^

### Characterization Analysis

The morphologies of HEK-293T-hACE2 NPs were observed by TEM (Tecnai G2 F20 U-TWIN, FEI, Hillsboro, OR, US). The DLS and ζ-potential experiments were determined by a Nano-ZS (Malvern, Worcestershire, UK) at room temperature. Protein contents were measured with a BCA kit (P0012, Beyotime). HEK-293T-hACE2 and HEK-293T NPs were denatured and resolved via 10% SDS-PAGE. Immunoblotting was applied to detect ACE2. The location of ACE2 on HEK-293T-hACE2 NPs was observed by immunofluorescence microscopy. The exact content of ACE2 in NPs was determined with a human ACE2 ELISA kit (ab235649, Abcam, Shanghai, CHN). ELISA was conducted in triplicate and repeated three times.

### Biolayer Interferometry (BLI)

The bindings of NPs to SARS-CoV-2 spike RBD (40592-V08H, Sino Biological) and D614G-S1 were measured using Forte Bio’s “Octet Red 96” BLI (Sartorius BioAnalytical Instruments, Bohemia, NY, US). Biotinylated SARS-CoV-2 RBD and D614G-S1 were obtained with a G-MM-IGT biotinylation kit (Genemore, Shanghai, CHN) was immobilized on SA biosensors at 15 μg mL^−1^. HEK-293T-hACE2 and HEK-293T NPs were prepared in sterile PBS. Association was performed at a shaking speed of 1000 rpm and ran for 300 s. To determine the affinity of ACE2 binding to D614G-S1, ACE2 was prepared in PBS with concentrations of 100, 75, 50, 25, and 10 nM. The running times for association and disassociation were both 300 s. The binding data were processed using Fortebio Data Analysis 7.0 software. The equilibrium dissociation constant (K_D_) and fit coefficient (R^2^) were generated by a 1:1 fitting model.

### Proteomics Research

HK-2 cells were exposed to 100 ng mL^−1^ of SARS-CoV-2 S1 at 37 °C for 48 h, in the absence and presence of 200 μg mL^−1^ of HEK-293T-hACE2 NPs. The total proteins were extracted and digested into peptides, which were then labeled with an amine-reactive TMTsixplex Isobaric Mass Tagging Kit (90064B, Thermo Fisher Scientific) and fractionated with reversed phase chromatography at pH 12 (Xbridge column, Waters, Shanghai, CHN) into 8 fractions. Peptides were resuspended in 0.1% formic acid (FA) and separated using U3000 RSLCnano liquid chromatography equiped with a Thermo Acclaim™ PepMap™ 100 C18 Trap Column (75 μm × 2 cm, 3 μm, 100 Å) and a C18a nalytical column (75 μm × 15 cm,, 3 μm, 100 Å). Elution was carried out at a flow rate of 400 nL min^−1^, in which solvent A (0.1% FA in water) and solvent B (0.1% FA in 80% acetonitrile) were applied. The elution gradient was 1-8% B (11 min), 8-30% B (99 min), 30-90% B (10 min), 90% B (2 min), 90-1% B (1 min), and 1% B (7 min).

The eluate was sprayed into an LTQ-Orbitrap Velos Pro MS spectrometer (Thermo Fisher Scientific) at a voltage of 2.5 kV. Full scan MS spectra (350-1800 m/z) were acquired at a resolution of 60,000. MS/MS spectra were acquired with a resolution of 30,000 by product ion scans (relative HCD energy 40) of the top 10 most abundant precursor ions in the survey scan. MS scans were recorded in profile mode, while the MS/MS was recorded in centroid mode. Three replicate injections were performed for each set of samples.

Data were processed using Proteome Discoverer software (PD) (version 1.4, Thermo Fisher Scientific). Peptides with scores above 20 and below the significance threshold filter (0.05) were selected for analysis. Single peptide identification required a score equal to or above the identity threshold. The MS/MS database searches were conducted using SEQUEST search algorithm through Proteome Discoverer platform (version 1.4, Thermo Fisher Scientific). The workflow included spectrum selector, SEQUEST search nodes, and percolator. Trypsin was specified as the protease and a maximum of two missed cleavages were allowed. MS and MS/MS mass tolerances were set to 10 ppm and 0.1 Da, respectively. False discovery rate of 1% was set at the PSM level as well as protein level. Gene names of encoded proteins identified in the proteomics analysis were uploaded into the online STRING database (Version 11.0) (https://string-db.org) for Gene Ontology (GO) annotation.

### Apoptosis Detection

The flow cytometry was performed as we recently discribed.^13^ Briefly, HK-2 was seeded in a 6-well plate at a density of 2×10^5^ CFU/well and cultured overnight in DMEM containing 10% FBS. Cells were co-incubated with 100 ng mL^−1^ of SARS-CoV-2 S1 for 12 h in the absence and presence of HEK-293T-hACE2 NPs (100 μg mL^−1^, based on the membrane protein). The cellular apoptosis was measured with a Beyotime Annexin V / PI detection kit (C1062M) and a BD flow cytometry system (Franklin Lakes, NJ, US), in which 15,000 events per sample were obtained. The experiment was conducted in triplicate and repeated twice, and the data were processed using FlowJo software (version 7.6.1).

### Pseudovirons Experiments

The adherence of SARS-CoV-2 S pseudovirons to HEK-293T-hACE2 NPs was observed with TEM, in which 40 μL of pseudovirons (1.98 × 10^7^ TU mL^−1^) were coincubated with 100 μL of HEK-293T-hACE2 NPs (2.5 mg mL^−1^, based on the concentration of membrane proteins) at 37 °C for 1 h. The luciferase assay was performed as we recently described.^8^ Briefly, HK-2 cells were seeded into a 96-well plate at a density of 5 × 10^3^ cells/well. SARS-CoV-2 S pseudovirons (20 μL, 1.98 × 10^7^ TU mL^−1^) pre-incubated with increasing concentrations of HEK-293T-hACE2 NPs (0.01-1 mg mL^−1^) at 37 °C for 1 h were added to HK-2 and incubated for 12 h. Cells were washed and lysed after 48 h of post-inoculation in DMEM containing 10% FBS. The luciferase activity was measured using a dual-luciferase reporter assay system (E1910, Promega, Beijing, CHN). The experiments were conducted in triplicate and repeated twice.

### Toxicological Evaluation

HUVECs obtained from the cell bank of CAS were seeded in a 96-well plate at a density of 5 × 10^3^ cells/well. HEK-293T-hACE2 NPs were prepared in DMEM containing 10% FBS with concentrations of 100, 200, 300, 400, and 500 μg mL^−1^. After adherence, cells were exposed to HEK-293T-hACE2 NPs at 37 °C for 24 h. The cell viability was measured using CCK-8 (Dojindo, Shanghai, CHN). Mice RBCs (300 μL) diluted in 0.9% NaCl solution were incubated with 1.2 mL of HEK-293T-hACE2 NPs at 37 °C for 2 h. The absorbance of the supernatant was determined at 405 nm using a microplate reader. Mice were cared for and treated in accordance with the National Institutes of Health (NIH) guidelines for the care and use of laboratory animals (NIH Publication No. 85e23 Rev. 1985) as approved by the Animal Experimental Ethics Committee of TMMU (AMUWEC2020799). The experiments were conducted in triplicate and repeated twice.

The toxicity of HEK-293T-hACE2 NPs *in vivo* was evaluated by a mouse experiment. HEK-293T-hACE2 NPs (25 mg kg^−1^) were administered by intravenous injection. Mouse blood obtained through the tail vein was analyzed on a Sysmex XT-2000i fully automatic hematology analyzer (Kobe, JPN) at 1, 3, and 5 days after injection. The mice were sacrificed on day 7. HE staining and an Olympus DX51 optical microscope (Tokyo, Japan) were applied to observe the pathological changes in major organs.

### Statistical Analysis

The significant difference (*P*) between each group was calculated using SPSS 16.0 software and the LSD multiple-comparison test. A *P* value lower than 0.05 was defined as statistically significant.

## ASSOCIATED CONTENT

Supporting Information

Figures and Tables are as noted in text.

## ACKNOWLEDGMENTS

This work was supported by grants from the National Natural Science Foundation of China (Nos. 81873605, 81725019, and 81703395), the program for scientific and technological innovation leader of Chongqing (CQYC201903084), the Natural Science Fund of Chongqing City (cstc2019jcyj-msxmX0011), and the frontier specific projects of Xinqiao Hospital (2018YQYLY004). There are no competing financial interests to declare. We would like to thank Prof. Lilin Ye (Institute of Immunology, PLA, Third Military Medical University) for instructing the pseudovirons experiment.

## Notes

### Competing Interest Statement

The authors have declared no competing interest.

