## Supplemental Table S1-S3, Figure S1-S4 for "Membrane Nanoparticles Derived from ACE2-rich Cells Block SARS-CoV-2 Infection"

**Contents:**

**Table S1**. Altered proteins of HK-2 cells exposed to SARS-CoV-2 S1.

**Table S2**. Altered signaling pathways of HK-2 cells exposed to SARS-CoV-2 S1.

**Table S3**. Altered proteins of HK-2 cells after exposure to SARS-CoV-2 S1 co-incubated with HEK-293T-hACE2 NPs.

**Figure S1**. Western blot determining the dose-dependent inhibition of HEK-293T-hACE2 NPs on SARS-CoV-2 S1.

**Figure S2**. Immunofluorescence microscopy revealing the inhibition of HD5 on SARS-CoV-2 S1 binding to HEK-293T-hACE2 NPs.

**Figure S3**. Online database showing the abundance of TMPRSS2 and furin in kidney.

**Figure S4**. Hemoglobin contents in mouse blood post-injection of HEK-293T-hACE2 NPs.

**Table S1**. Altered proteins of HK-2 cells exposed to SARS-CoV-2 S1.

| **Accession** | **Annotation** | **Contents relative to the control** | ***P*-value** |
| --- | --- | --- | --- |
| **Down-regulation** |  |  |  |
| **Q8WXE9** | Stonin-2 | 0.643333603 | 0.021942109 |
| **Q5T3I0** | G patch domain-containing protein 4 | 0.678983566 | 0.034658609 |
| **Q8IXU6** | Solute carrier family 35 member F2 | 0.706658319 | 0.04184812 |
| **Q8N4Q1** | Mitochondrial intermembrane space import and assembly protein 40 | 0.736118658 | 0.04175746 |
| **A4D1E9** | GTP-binding protein 10 | 0.760237446 | 0.002666624 |
| **P04264** | Keratin, type II cytoskeletal 1 | 0.763221681 | 0.014976794 |
| **Q9NW81** | Distal membrane-arm assembly complex protein 2 | 0.767410469 | 0.00854136 |
| **Q5XKP0** | MICOS complex subunit MIC13 | 0.769301966 | 0.021141438 |
| **P55085** | Proteinase-activated receptor 2 | 0.77271055 | 0.044738402 |
| **Q8IYT3** | Coiled-coil domain-containing protein 170 | 0.777114384 | 0.017082045 |
| **Q6NUQ1** | RAD50-interacting protein 1 | 0.780624975 | 0.045856989 |
| **O60507** | Protein-tyrosine sulfotransferase 1 | 0.787835705 | 0.027159869 |
| **P82664** | 28S ribosomal protein S10, mitochondrial | 0.798344862 | 0.036848549 |
| **P20337** | Ras-related protein Rab-3B | 0.809207744 | 0.036058712 |
| **Q96FN4** | Copine-2 | 0.814906151 | 0.031981358 |
| **O00767** | Acyl-CoA desaturase | 0.816080912 | 0.037271623 |
| **Q8IYB8** | ATP-dependent RNA helicase SUPV3L1, mitochondrial | 0.817639987 | 0.028221344 |
| **P0CG12** | Decreased expression in renal and prostate cancer protein | 0.818224936 | 0.002038395 |
| **Q9BV81** | ER membrane protein complex subunit 6 | 0.818947736 | 0.005229445 |
| **P11233** | Ras-related protein Ral-A | 0.823603525 | 0.001912781 |
| **Q01970** | 1-phosphatidylinositol 4,5-bisphosphate phosphodiesterase beta-3 | 0.824142378 | 0.00595548 |
| **P30405** | Peptidyl-prolyl cis-trans isomerase F, mitochondrial | 0.827273895 | 0.019031608 |
| **Q13277** | Syntaxin-3 | 0.830354388 | 0.016268929 |
| **Up-regulation** |  |  |  |
| **P61604** | 10 kDa heat shock protein, mitochondrial | 0.835272223 | 0.00378986 |
| **P05141** | ADP/ATP translocase 2 | 0.835769919 | 0.019017606 |
| **Q9UHA4** | Ragulator complex protein LAMTOR3 | 0.836943661 | 0.033954244 |
| **O75323** | Protein NipSnap homolog 2 | 0.842229108 | 0.042302425 |
| **Q14197** | Peptidyl-tRNA hydrolase ICT1, mitochondrial | 0.844100719 | 0.037710269 |
| **Q969P6** | DNA topoisomerase I, mitochondrial | 0.845421353 | 0.007641381 |
| **Q92930** | Ras-related protein Rab-8B | 0.846483571 | 0.011314035 |
| **Q96A57** | Transmembrane protein 230 | 0.848965511 | 0.022146857 |
| **Q86VM9** | Zinc finger CCCH domain-containing protein 18 | 0.849116629 | 0.003751821 |
| **P0DJ07** | Protein PET100 homolog, mitochondrial | 0.84930686 | 0.012755015 |
| **P09622** | Dihydrolipoyl dehydrogenase, mitochondrial | 0.849738025 | 0.003046366 |
| **Q9Y239** | Nucleotide-binding oligomerization domain-containing protein 1 | 0.852739155 | 0.030104383 |
| **P40926** | Malate dehydrogenase, mitochondrial | 0.855317881 | 0.001298213 |
| **Q96AG4** | Leucine-rich repeat-containing protein 59 | 0.855632909 | 0.007199102 |
| **Q9UDW1** | Cytochrome b-c1 complex subunit 9 | 0.856964651 | 0.018348993 |
| **Q9BYG3** | MKI67 FHA domain-interacting nucleolar phosphoprotein | 0.857566016 | 0.014917217 |
| **P53801** | Pituitary tumor-transforming gene 1 protein-interacting protein | 0.858766486 | 0.042678502 |
| **Q08722** | Leukocyte surface antigen CD47 | 0.859383449 | 0.025358001 |
| **Q15084** | Protein disulfide-isomerase A6 | 0.859750008 | 0.014165478 |
| **Q15392** | Delta(24)-sterol reductase | 0.860096564 | 0.019351718 |
| **P24390** | ER lumen protein-retaining receptor 1 | 0.860692737 | 0.038062906 |
| **P51571** | Translocon-associated protein subunit delta | 0.860813636 | 0.013473939 |
| **P99999** | Cytochrome c | 0.861221536 | 0.008489706 |
| **P55145** | Mesencephalic astrocyte-derived neurotrophic factor | 0.86195893 | 0.00359699 |
| **P84157** | Matrix-remodeling-associated protein 7 | 0.862367809 | 0.013983643 |
| **Q6ZRP7** | Sulfhydryl oxidase 2 | 0.862569896 | 0.03397915 |
| **Q9BQC6** | Ribosomal protein 63, mitochondrial | 0.86283035 | 0.01609103 |
| **O00483** | Cytochrome c oxidase subunit NDUFA4 | 0.863050397 | 0.013349224 |
| **Q99643** | Succinate dehydrogenase cytochrome b560 subunit, mitochondrial | 0.863095017 | 0.025129908 |

**Table S2**. Altered signaling pathways of HK-2 cells exposed to SARS-CoV-2 S1.

| **Signaling pathways** | **Observed gene count** | **Matched proteins** | **False discovery rate** |
| --- | --- | --- | --- |
| Calcium signaling pathway | 5 | PLCB3,PPIF,VDAC1,VDAC2,VDAC3 | 0.0049 |
| cGMP-PKG signaling pathway | 5 | PLCB3,PPIF,VDAC1,VDAC2,VDAC3 | 0.0049 |
| NOD-like receptor signaling pathway | 5 | FADD,PLCB3,VDAC1,VDAC2,VDAC3 | 0.0049 |
| Cholesterol metabolism | 3 | VDAC1,VDAC2,VDAC3 | 0.0051 |
| Necroptosis | 4 | FADD,VDAC1,VDAC2,VDAC3 | 0.0084 |
| Ferroptosis | 2 | VDAC2,VDAC3 | 0.0455 |

**Table S3**. Altered proteins of HK-2 cells after exposure to SARS-CoV-2 S1 co-incubated with HEK-293T-hACE2 NPs.

| **Accession** | **Annotation** | **Contents relative to the control** | ***P*-value** |
| --- | --- | --- | --- |
| **Down-regulation** |  |  |  |
| P15408 | Fos-related antigen 2 | 0.601894698 | 0.011595477 |
| Q9NW81 | Distal membrane-arm assembly complex protein 2 | 0.656528382 | 0.008068172 |
| P55085 | Proteinase-activated receptor 2 | 0.683834421 | 0.044202837 |
| P02792 | Ferritin light chain | 0.784046364 | 0.032938325 |
| Q92504 | Zinc transporter SLC39A7 | 0.79531332 | 0.002421933 |
| P07477 | Trypsin-1 | 0.809153345 | 0.031506015 |
| Q8IYT3 | Coiled-coil domain-containing protein 170 | 0.820099066 | 0.008295192 |
| Q99618 | Cell division cycle-associated protein 3 | 0.821492591 | 0.024032876 |
| Q5XKP0 | MICOS complex subunit MIC13 | 0.828911819 | 0.00152105 |
| Q9Y6X3 | MAU2 chromatid cohesion factor homolog | 0.832943672 | 0.00462652 |
| **Up-regulation** |  |  |  |
| Q8TBX8 | Phosphatidylinositol 5-phosphate 4-kinase type-2 gamma | 0.836946955 | 0.033441818 |
| Q8TDQ7 | Glucosamine-6-phosphate isomerase 2 | 0.845329971 | 0.013033796 |
| Q13907 | Isopentenyl-diphosphate Delta-isomerase 1 | 0.857596926 | 0.034245908 |
| Q01970 | 1-phosphatidylinositol 4,5-bisphosphate phosphodiesterase beta-3 | 0.858027544 | 0.016093966 |
| P20936 | Ras GTPase-activating protein 1 | 0.864356926 | 0.014135582 |
| O14817 | Tetraspanin-4 | 0.867482572 | 0.032484706 |
| P62861 | 40S ribosomal protein S30 | 0.867658419 | 0.004683421 |
| Q8NF91 | Nesprin-1 | 0.871888303 | 0.022955789 |
| Q9UBV2 | Protein sel-1 homolog 1 | 0.874340821 | 0.017047553 |
| P50542 | Peroxisomal targeting signal 1 receptor | 0.877882517 | 0.00555954 |
| Q9BRX8 | Peroxiredoxin-like 2A | 0.878993655 | 0.042129291 |
| P34896 | Serine hydroxymethyltransferase, cytosolic | 0.879344941 | 0.033635415 |
| O95249 | Golgi SNAP receptor complex member 1 | 0.879374717 | 0.016912753 |
| Q9UQ90 | Paraplegin | 0.884290277 | 0.047794451 |
| Q8IYB3 | Serine/arginine repetitive matrix protein 1 | 0.884777857 | 0.018250166 |
| Q7Z6E9 | E3 ubiquitin-protein ligase RBBP6 | 0.88495513 | 0.011380209 |
| Q13637 | Ras-related protein Rab-32 | 0.885927257 | 0.045245456 |
| Q13601 | KRR1 small subunit processome component homolog | 0.887129469 | 0.019060449 |
| P84157 | Matrix-remodeling-associated protein 7 | 0.887141693 | 0.038304286 |
| P11166 | Solute carrier family 2, facilitated glucose transporter member 1 | 0.887693819 | 0.016047611 |
| P12236 | ADP/ATP translocase 3 | 0.888624973 | 0.020236036 |

**Figure S1**. Western blot determining the dose-dependent inhibition of HEK-293T-hACE2 NPs on SARS-CoV-2 S1. HK-2 cells were exposed to 20 μg mL^-1^ of His-tag-containing SARS-CoV-2 S1 in the absence and presence of HEK-293T-hACE2 NPs. A primary anti-His-tag mouse monoclonal antibody and a goat anti-mouse secondary antibody were employed to detect SARS-CoV-2 S1. β-actin is the reference.

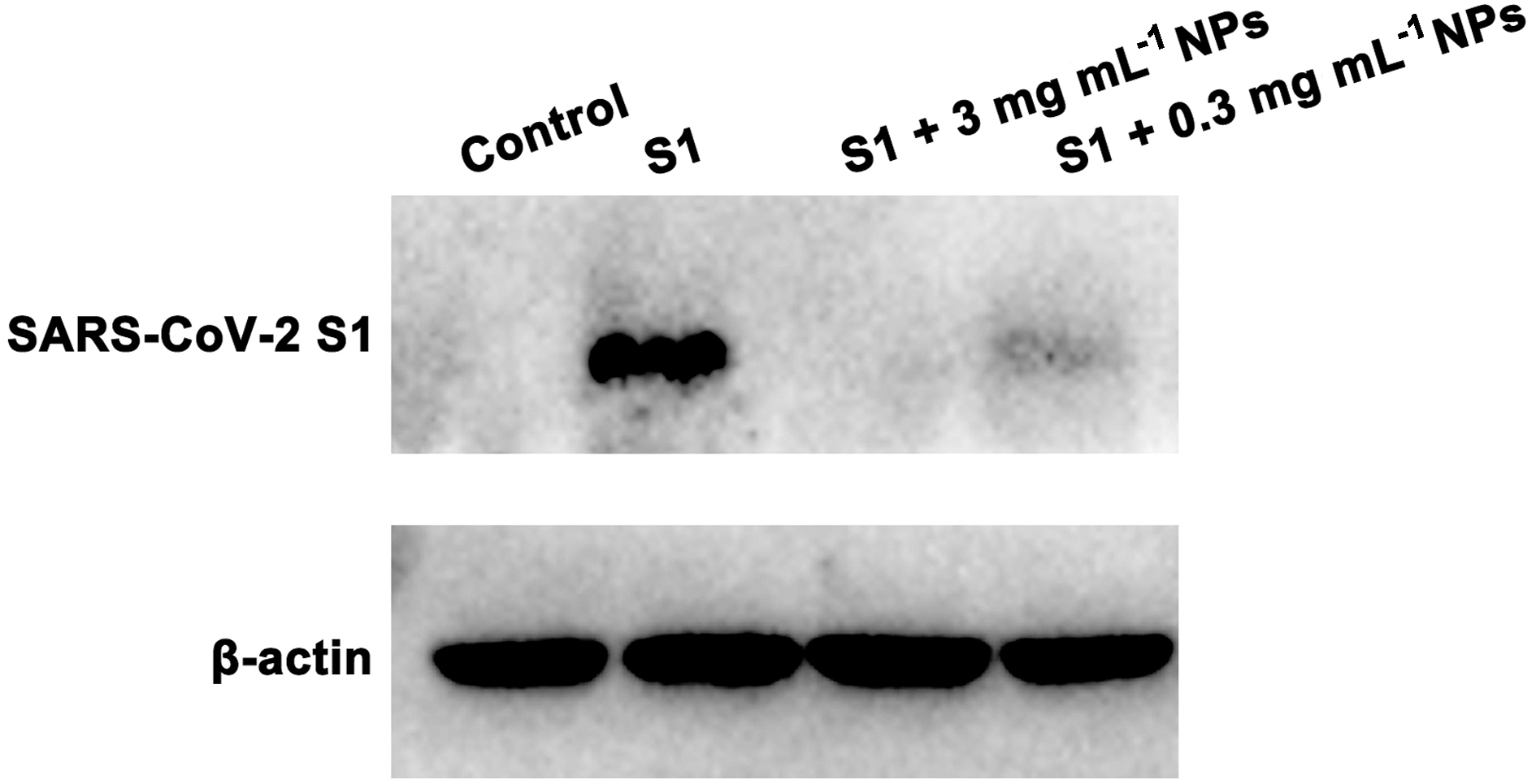

**Figure S2**. Immunofluorescence microscopy revealing the inhibition of HD5 on SARS-CoV-2 S1 binding to HEK-293T-hACE2 NPs. HEK-293T-hACE2 NPs cloaked by 100 μg mL^-1^ of HD5 were co-incubated with 10 μg mL^-1^ of SARS-CoV-2 S1 at 37 °C for 1 h. A primary anti-spike rabbit monoclonal antibody and a goat anti-rabbit secondary antibody (Alexa Fluor 488) were used to stain S1. Scale bar indicates 20 μm.

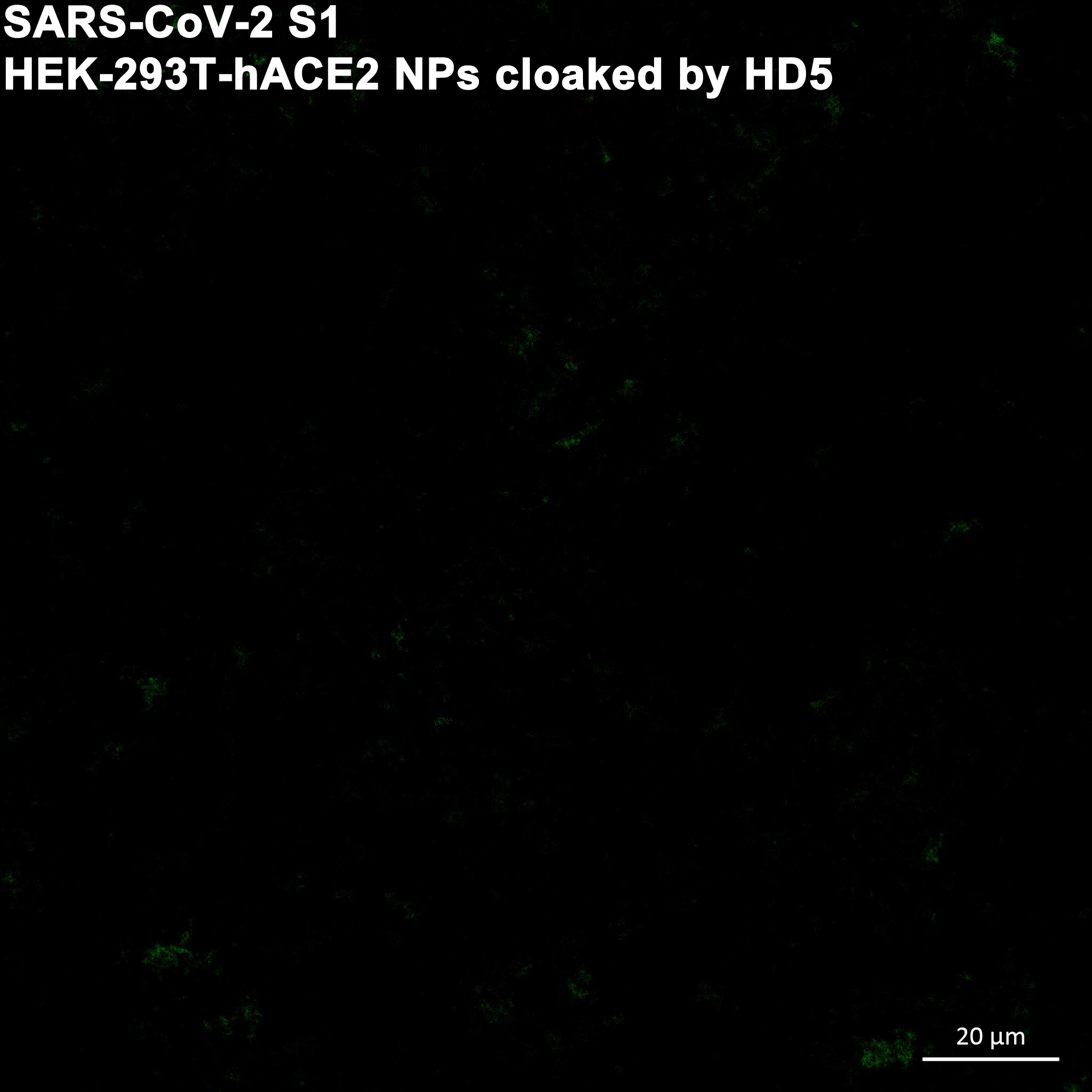

**Figure S3**. Online database showing the abundance of TMPRSS2 and furin in kidney. The contents of TMPRSS2 (a) and furin (b) *in vivo*, as revealed by The Human Protein Atlas (http://www.proteinatlas.org/). Kidney and urinary bladder are highlighted in red frames.

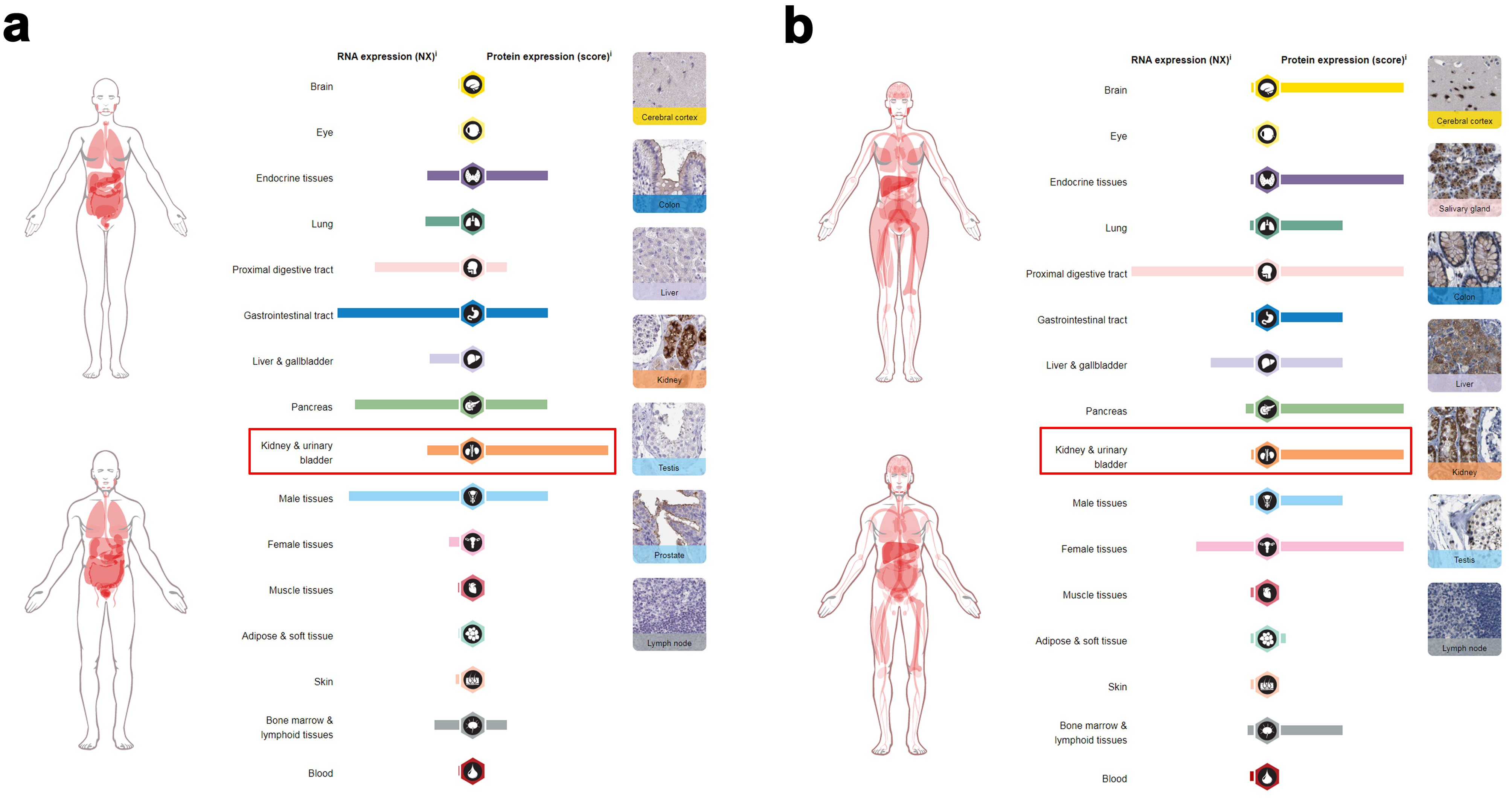

**Figure S4**. Hemoglobin contents in mouse blood post-injection of HEK-293T-hACE2 NPs. HEK-293T-hACE2 NPs are intravenous administrated at 25 mg kg^-1^. Shown are the counts of hemoglobin (HGB) in mouse blood at different days. The results are presented as the means ± SD.

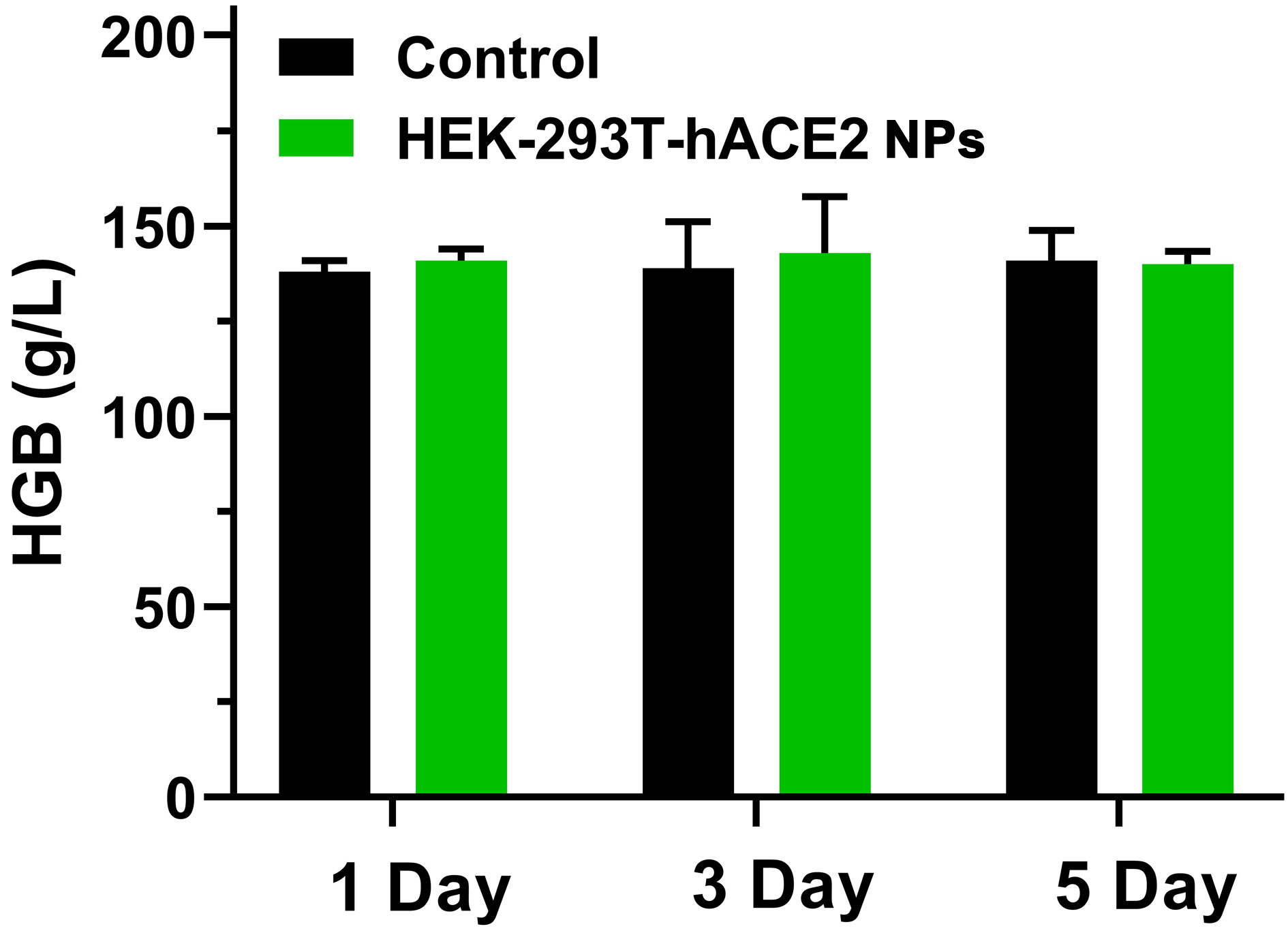
